# Calcium waves facilitate and coordinate the contraction of endfeet actin stress fibers in *Drosophila* interommatidial cells

**DOI:** 10.1101/2021.04.08.439074

**Authors:** Donald F. Ready, Henry C. Chang

## Abstract

Actomyosin contraction shapes the *Drosophila* eye’s panoramic view. The convex curvature of the retinal epithelium, organized in ∼800 close-packed ommatidia, depends upon a fourfold condensation of the retinal floor mediated by contraction of actin stress fibers in the endfeet of inter<u>o</u>mmatidial <u>c</u>ells (IOCs). How these tensile forces are coordinated is not known. Here, we discover a novel phenomenon: Ca^2+^ waves regularly propagate across the IOC network in pupal and adult eyes. Genetic evidence demonstrates that IOC Ca^2+^ waves are independent of phototransduction, but require inositol 1,4,5-triphosphate receptor (IP3R), suggesting these waves are mediated by Ca^2+^ releases from ER stores. Removal of *IP3R* disrupts stress fibers in IOC endfeet and increases the basal retinal surface by ∼40%, linking IOC waves to facilitating stress fiber contraction and floor morphogenesis. Further, *IP3R* loss disrupts the organization of a collagen IV network underneath the IOC endfeet, implicating ECM and its interaction with stress fibers in eye morphogenesis. We propose that coordinated Ca^2+^ spikes in IOC waves promote stress fiber contractions, ensuring an organized application of the planar tensile forces that condense the retinal floor.

**Summary Statement:** Ca^2+^ waves have an important role in generating tensile forces to shape the *Drosophila* eye’s convex curvature. Coordinated Ca^2+^ spikes facilitate actomyosin contractions at the basal endfeet of interommatidial cells.

## Introduction

Tissue construction during development often requires a specific subset of its constituent cells to undergo orchestrated shape change. To accomplish this, tensile forces from actomyosin networks, which are critical in maintaining and regulating cell shapes, need to be spatially and temporally coordinated in a selective cell population (Guillot and Lecuit, 2013, Paluch and Heisenberg, 2009, Heisenberg and Bellaiche, 2013). While the mechanism behind the actomyosin contraction in vitro and in individual cells has been extensively studied, how these forces are coordinated across a cell field during tissue formation is not well understood.

To investigate the mechanisms coordinating contractile forces critical for tissue morphogenesis, we have used the *Drosophila* compound eye, a convex epithelium with stereotypical cell composition and arrangement (Waddington, 1962). The compound eye consists of ∼800 unit eyes, ommatidia, each comprising a distal corneal lens secreted by four cells and two primary pigment cells positioned above eight photoreceptors elongated on the optical axis. Ommatidia are embedded in a honeycomb lattice of inter<u>o</u>mmatidial <u>c</u>ells (IOCs), pigment-secreting cells that separate and optically insulate ommatidia throughout. At the retinal floor, IOCs flatten their endfeet to form the fenestrated membrane, an ECM/cytoskeletal specialization that defines the proximal surface of the eye (Fig. 1). This reiterative nature of the eye architecture implies the tensile forces for cell morphogenesis in one cluster are similarly applied in all other ommatidia across the retina, making the eye uniquely suited for analyzing how local forces are coordinated to ensure proper tissue formation.

**Figure 1.**
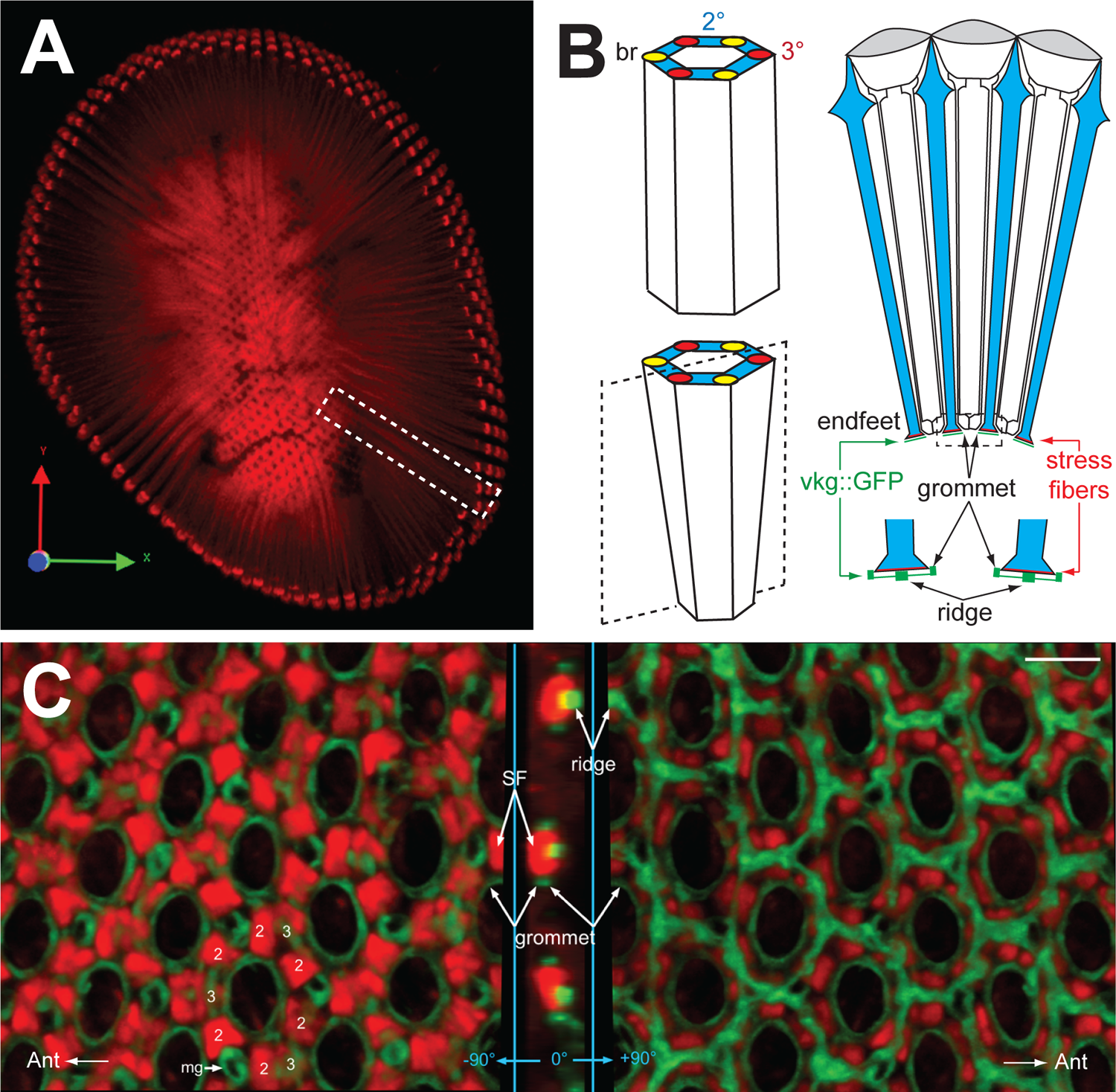
*Drosophila* eye curvature requires retinal floor condensation. (A) A snapshot of 3D rendering from confocal images of an adult retina stained with phalloidin. The projection reveals the rhabdomeres extended throughout the depth of this convex epithelium (dashed box) and the endfeet stress fibers at the retinal floor (center). (B) A schematic diagram showing a hexagonal column, consisted of secondary (blue), tertiary (red) pigment cells, and bristle cells (yellow), with and without basal reduction. A cross-section drawing (angle of the section indicated with a dashed box) showing the secondary pigment cell (blue) endfeet of three adjacent clusters. IOC endfeet, stand upon planar basement membrane ECM, organized in grommets and ridges (green); Actin stress fibers (red lines) within the endfeet are anchored to adjacent grommets. Corneal lenslets are indicated with gray ovals. (C) A floor triptych rendered from confocal stacks of *cn, bw, vkg::GFP* adult retina, stained with phalloidin (red). Left: a view from within the retina, showing the stereotypical arrangement of dense stress fiber patches (SF) in the endfeet of secondary and tertiary pigment cells. Bristle axons exit the retina via mini grommets (mg). Middle: the reconstruction edge-on view at the indicated cut. Right: a bottom view revealing retinal ECM organized in grommets and ridges. Ridges join to form a hexagonal network. Scale bar=5μm.

Actomyosin-dependent contraction has been implicated in several processes during fly eye development, including the apical cell constriction in the morphogenetic furrow (MF; an indentation that sweeps across the larval eye disc) (Benlali et al., 2000, Fernandes et al., 2014), the pigment cell shape and photoreceptor position in the epithelium (Warner and Longmore, 2009a, Warner and Longmore, 2009b, Lee and Treisman, 2004), rhabdomere biogenesis (Baumann, 2004, Galy et al., 2011), and the retinal lumen formation (Nie et al., 2014). In addition, actomyosin tensile forces participate in the reduction of retinal basement, a process critical for the formation of eye’s curvature. During the latter half of pupal development, the IOC endfeet undergo a fourfold contraction (Longley and Ready, 1995), shaping the ommatidia into elongated hexagonal frustums that pack to give the eye its panoramic convex curvature (Fig. 1). Planar tension at the retinal floor has the additional role of aligning each ommatidium’s rhabdomeres, photosensory waveguides, along the optical axis. Rhabdomeres are sprung between the rigid outer dome of the faceted cornea and the inner retinal floor in a cage of specialized cell-cell junctions (Cagan and Ready, 1989) (Fig. 1). Tension in the convex retinal floor stretches the rhabdomere cage on the longitudinal axis, which aligns rhabdomeres and must be balanced with growth and renewal of rhabdomeres across the retina (Raghu et al., 2009).

The reduction in basal surface is largely powered by the actin stress fibers and the associated non-muscle myosin II in IOC endfeet, as eyes deficient in *zipper* (*zip*; myosin II heavy chain) and actin regulation contain distorted retinal floors (Baumann, 2004, Cagan and Ready, 1989, Galy et al., 2011, Longley and Ready, 1995). At the mid-pupal stage, the stress fibers form at the IOC endfeet and are anchored to the grommets via integrins (Longley and Ready, 1995), appearing as filaments emanated from the grommets. These stress fibers mature through the late pupal stages and generate tension, bringing attached grommets closer together. Concomitant with contraction, stress fibers transform from a looser, filamentous organization to larger dense patches with higher levels of stress fibers abutting the neighboring grommets (partially contracted), and then to small dense patches connecting the grommets (fully contracted). Thus, the contraction of endfeet stress fibers, manifested by a progressive reduction of their profiles, compacts the fenestrated membrane, including the ECM below. The regular organization of the ommatidia is preserved at the retinal floor, suggesting that the tensile forces for reducing the IOC endfeet are precisely applied.

To decipher the mechanism coordinating IOC endfeet contraction, we have combined live imaging with a Ca^2+^ sensor (Chen et al., 2013) and mutational analysis to investigate Ca^2+^ signaling in non-neuronal retinal cells. Ca^2+^ is a ubiquitous and versatile second messenger, impacting virtually all cell physiologies (Berridge et al., 2000). In addition to facilitating muscle contraction, Ca^2+^ signaling participates in actin remodeling and contraction in non-muscle cells. Ca^2+^-dependent regulation of non-muscle myosin II, a hexamer of two heavy chains, two regulatory light chains (MRLC), and two essential light chains, has been well characterized (Brito and Sousa, 2020, Heissler and Sellers, 2016). Myosin II activity is increased by the phosphorylation of MRLC, which can be upregulated by calmodulin-dependent myosin light chain kinase (MLCK) in response to Ca^2+^ signaling (Scholey et al., 1980, Heissler and Sellers, 2016).

Thus, an orchestrated Ca^2+^ signaling, like a propagation of Ca^2+^ spikes through constituent cells, is capable of coordinating the activity of actomyosin networks in tissue morphogenesis (Homolya et al., 2000). In *Drosophila* egg activation, a single Ca^2+^ wave leads to actin reorganization and the formation of dynamic actin around cell cortex (York-Andersen et al., 2020, Kaneuchi et al., 2015, York-Andersen et al., 2015). In wing discs, oscillating Ca^2+^ waves influence actomyosin organization (Balaji et al., 2017) and facilitate recovery in response to mechano-injury (Brodskiy et al., 2019, Narciso et al., 2017, Restrepo and Basler, 2016). In vertebrate epithelia, Ca^2+^ propagation induces actin rearrangement in the surrounding cells to promote cell extrusion (Takeuchi et al., 2020).

In mammalian retina, Ca^2+^ signaling is known to facilitate the formation of electrical synapses and mediates trophism between supporting cells and neurons (Barres, 2008, Allaman et al., 2011, Halassa and Haydon, 2010, Fields and Stevens-Graham, 2002). In addition, intercellular propagations of Ca^2+^ spikes sweep across glial cell networks, including CNS astrocytes (Cornell-Bell et al., 1990) and Müller retinal glial cells (Kurth-Nelson et al., 2009), although the roles of Ca^2+^ waves in these cells remain unresolved (Scemes and Giaume, 2006). In *Drosophila* retina, Ca^2+^ has critical functions in phototransduction (Voolstra and Huber, 2020, O’Tousa, 2002) and Ca^2+^ dynamics in isolated live photoreceptors have been recorded (Asteriti et al., 2017, Hardie, 1996). Stimulus-independent Ca^2+^ oscillations have also been observed in developing photoreceptors and are thought to facilitate neuronal connections (Akin et al., 2019). The versatility of Ca^2+^ signaling and its known impact on actomyosin activity make it a good candidate for orchestrating endfeet stress fiber contraction, although whether Ca^2+^ signaling has a role in accessory cells in fly retina is not known.

Here we report a novel observation: Ca^2+^ waves regularly sweep across the eye’s honeycomb-like lattice of IOCs. These waves, formed during the pupal development, require IP3R, indicating that IOC Ca^2+^ spikes are mediated by Ca^2+^ release from internal stores. Mutational analysis of *IP3R* suggests that Ca^2+^ waves contribute to the retinal floor reduction by facilitating the contraction of actin stress fibers at the IOC endfeet. Our work suggests a mechanism by which intercellular Ca^2+^ waves coordinate contractile forces for an orderly retinal floor condensation. Moreover, *IP3R^-^* deforms fenestrated membrane ECM, indicating that contractile forces from the endfeet stress fibers shape the ECM network.

## Results

### A model for visualizing intracellular Ca^2+^ levels in *Drosophila* retina

To monitor the intracellular Ca^2+^ levels in developing and adult *Drosophila* eyes, we expressed GCaMP6m (Chen et al., 2013) with *longGMR-GAL4* (Wernet et al., 2003), hereafter referred to as *lGMR>GCaMP6m*. Like the more widely used *GMR-GAL4* (Freeman, 1996), *lGMR-GAL4* becomes active in cells posterior to the MF and remains active in most of the retinal cells at subsequent stages (except the bristle cells). We chose *lGMR-GAL4* because its moderate GAL4 expression level, as compared to *GMR-GAL4*, is less likely to cause cytotoxicity, which may damage the eye architecture and indirectly perturb the cellular Ca^2+^ level. To minimize potential toxicity and artifact from high GAL4 expression, all *lGMR>GCaMP6m* recordings presented in subsequent sections were performed with heterozygotes.

Using this *lGMR>GCaMP6m* platform, we analyzed the spatiotemporal pattern of cytosolic Ca^2+^ increases in non-neuronal supporting cells, namely IOCs, primary pigment cells, and cone cells. As the composition, arrangement, and architecture of these accessory cells are well established (Cagan and Ready, 1989), we imaged *lGMR>GCaMP6m* retinas at different focal planes to monitor their respective Ca^2+^ activities.

### Ca^2+^ waves propagate through interommatidial cells

Imaging young *lGMR>GCaMP6m* adult retina at the distal plane showed Ca^2+^ spikes propagated regularly across the hexagonal IOC lattice (Fig. 2A, Movie 1; hereafter referred to as IOC waves). We tracked the movement of the wave fronts by sequentially subtracting signal intensities from consecutive images of the time-lapsed series. This operation revealed that the initiation of IOC waves was not restricted to a specific region. For instance, the IOC waves in Movie 1 could initiate from the middle (t=80s), anterior (t=4s, 152s, 210s, 266s), anterior-dorsal (t=92s), posterior-ventral (t=14s), and posterior-dorsal (t=20s) edges of this *lGMR>GCaMP6m* retina (Fig. 2B). Once initiated, the IOC waves moved equally in all directions across the eye field (Fig. 2C). Each IOC spiked approximately 2s after a neighboring cell, translating to a wave speed of 4-5um/s. When two IOC waves collided, the wave fronts merged and advanced toward regions where the cells had yet to experience recent Ca^2+^ spikes (Fig. 2D). If no such region was available, the colliding IOC waves ceased (Fig. 2D). The GCaMP6m intensity at the juncture of merging wave fronts was not greater than those before the collision, implying that one or more factors responsible for triggering IOC Ca^2+^ spikes are limiting, rendering these cells unresponsive to additional dose of spike-inducing signal.

**Figure 2.**
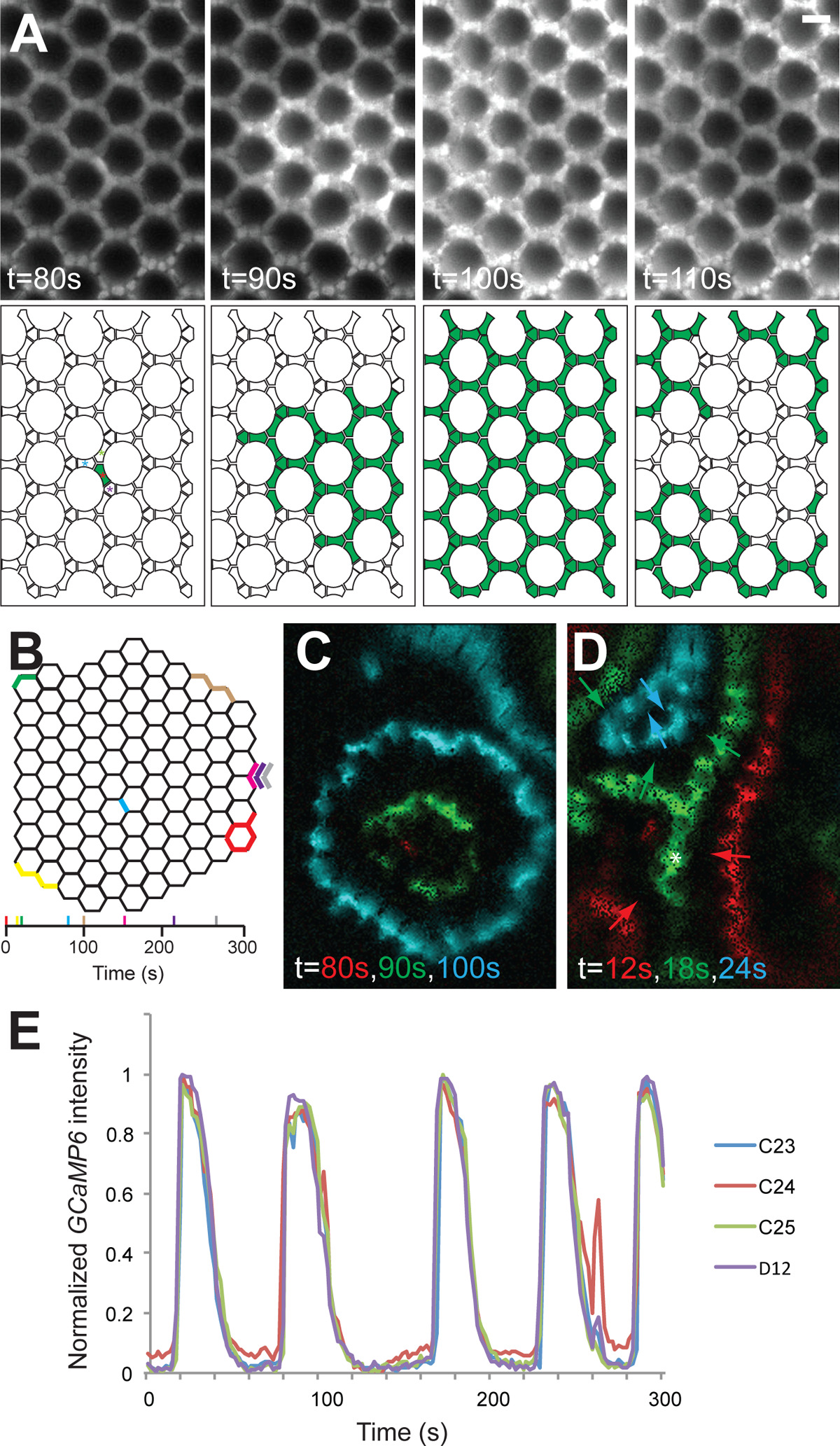
Ca^2+^ waves propagate regularly through IOC cells. (A) Top row: a montage showing a Ca^2+^ wave initiated at t=80s in a 1-day-old *lGMR>GCaMP6m* retina. Bottom row: schematic drawings of the above micrographs, showing the IOCs experiencing Ca^2+^ spikes (green). Bristle cells are omitted from the drawings. Scale bar=10μm. (B) A schematic representation of the IOC wave origins during the 300s recording. For those IOC waves moving from the edges, the exact origin cells cannot be determined and the first cells with Ca^2+^ spikes when these waves come into view are indicated by colors. The color code for each event and its corresponding time is shown in the line below. (C) An image overlaying three wave fronts (generated by intensity subtraction of consecutive images) at t=80s (red), 90s (green), and 100s (blue) respectively. This IOC wave (the same one in A and corresponds to the light blue event in B), initiated at t=80s, propagates in all directions at 5μm/s. At t=100s, a distinct wave front (the brown event in B) approaches from the anterior-dorsal edge. (D) An image overlaying three wave fronts at t=12s (red), 18s (green), and 24s (blue) respectively (the wave directions are indicated by arrows). At t=12s, two wave fronts approach from anterior and posterior-ventral edges. At t=18s, these two wave fronts merge into one (the merging site is indicated by an asterisk) and move dorsally, and a distinct wave approaches from the posterior-dorsal edge. At t=24s, all these waves merged to form a centripetal front. (E) Plot of normalized *GCaMP6m* intensity (*(F-F_min_)/(F_max_-F_min_)*) over time showing the regular Ca^2+^ oscillation in 4 cells, labeled in the first panel of schematic drawing in A.

The characteristics of Ca^2+^ spikes within these IOC waves were remarkably consistent (Fig. 2E). In Movie 1, all 273 cells (204 secondary and 69 tertiary pigment cells) exhibited regular calcium oscillations (average peak number was 4.8 ± 0.4, n=273) during the 5-minute recording. These reproducible Ca^2+^ spikes rose sharply to a maximum in approximately 6.8 ± 2.2s (Ton; time from baseline to peak; n=1309 peaks) with a slower return to baseline (25.1 ± 3.8s; Toff; time from peak to baseline). The interval time, measured as the duration from one peak to the subsequent peak, was 68.6 ± 12.4s. The peak intensities of Ca^2+^ spikes experienced by individual cells showed no apparent dampening over time. No noticeable difference in Ca^2+^ spike characteristics was seen between secondary and tertiary pigment cells (Fig. S1B) or between cells from different regions (Fig. S1C). Pair-wise Pearson correlation analysis performed on the numeric series generated from GCaMP6m intensity over time indicated that the cells were most related to their adjacent cells, further confirming that the wave-like nature of these Ca^2+^ spikes (Fig. S1D). To demonstrate these descriptions are representative, we present the analysis of IOC waves (still images shown in Fig. 5D) from an independently acquired *lGMR>GCaMP6m* recording in Fig. S2 (n=474 cells).

### IOC waves include primary pigment cells, but not the cone cells

To determine whether other accessory cells contribute to IOC waves, we imaged young *lGMR>GCaMP6m* retinas more proximally, at which Ca^2+^ increases in IOC waves, while not being easily attributed to individual cells, were still recognizable (Fig. 3A). At this plane, *GCaMP6m* signals in the nuclei of primary pigment cells, the two crescent-shaped cells situated apically between the IOC and cone cells, were particularly noticeable, as the pigment granules were thought to quench the *GCaMP6m* fluorescence in the cytoplasm (Fig. 3A). From the two independent recordings (Movies 2 and 3), *GCaMP6m* signals in the primary pigment cells increased synchronously with IOC wave passages (Fig. 3A and B, arrows; Fig. S3A and B), demonstrating that the IOC wave network includes the primary pigment cells. In support of this, the characteristics of Ca^2+^ spikes in primary pigment cells resembled to those observed in the secondary and tertiary pigment cells (Fig. 3C and Fig. S3C).

**Figure 3.**
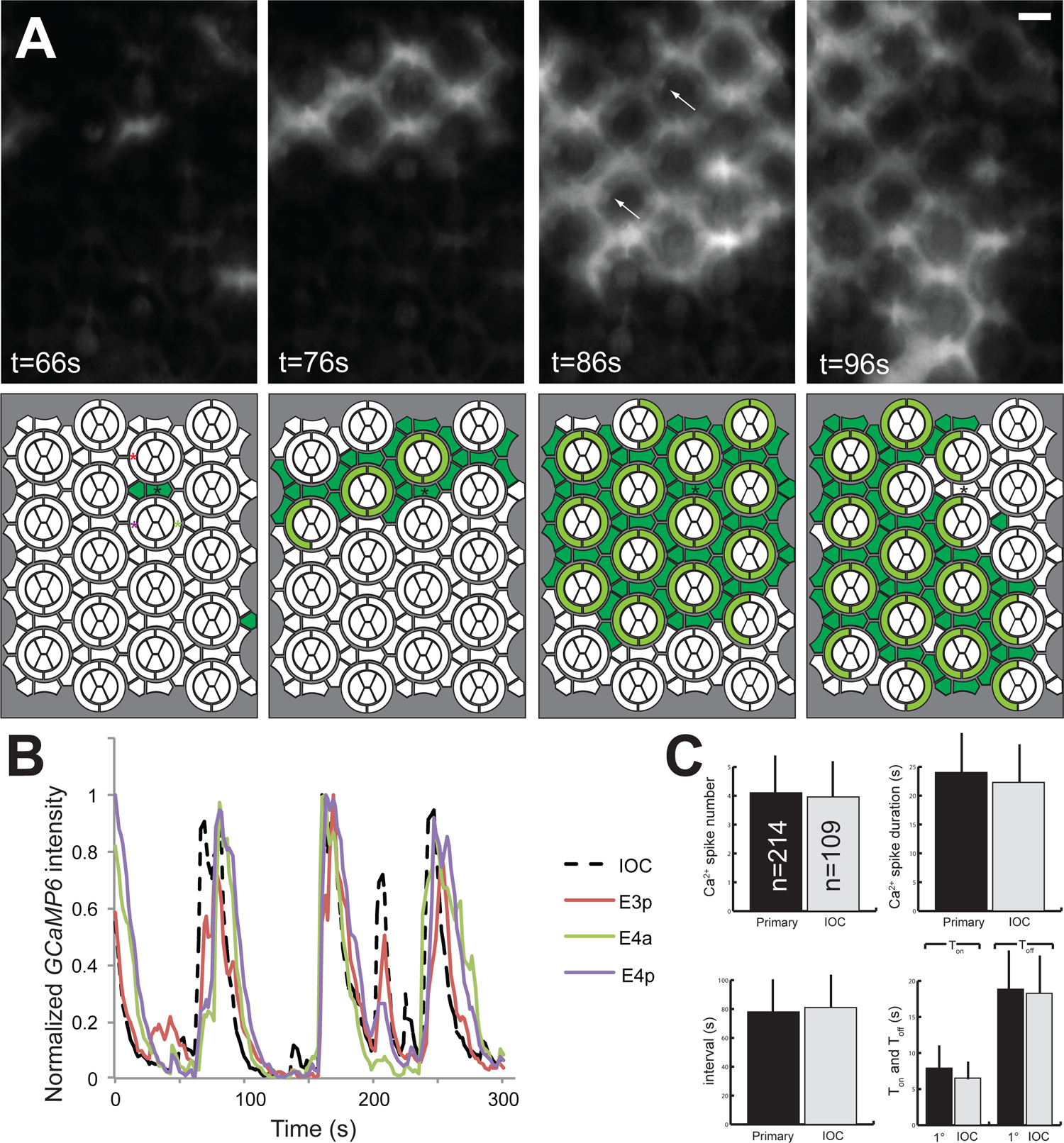
primary pigment cells, but not cone cells, are a part of the IOC wave network. (A) Top row: a young *lGMR>GCaMP6m* retina imaged at a proximal plane, showing an IOC wave initiated from a secondary pigment cell (asterisk in the drawing) at t=66s traversing through the primary pigment cells. In primary pigment cells, the *GCaMP6m* signals in the nuclei are stronger than those seen in the cytoplasm (arrows). Bottom row: schematic drawings showing the 4 pie-slice-shaped cone cells, the two crescent-shaped primary pigment cells, and the interommatidial secondary and tertiary pigment cells. IOC and primary pigment cells experiencing Ca^2+^ spikes are highlighted in green and olive green respectively. As GCaMP6m signals from secondary and tertiary pigment cells at this plane are hazy, those assigned to exhibit Ca^2+^ spikes are approximations. Scale bar=10μm. (B) Plot of normalized *GCaMP6m* intensity over time showing the Ca^2+^ oscillation in one secondary (dashed black line; from the cell marked with black asterisk) and three primary (solid color lines; from cells labeled with corresponding color asterisks) pigment cells. (C) Comparison of Ca^2+^ spike characteristics between IOC (n=109) and primary pigment cells (n=214) from this recording.

In contrast, cone cells, the four cells situated centrally atop the photoreceptors, exhibited Ca^2+^ activities independent of the IOC waves (Fig. 4A, Movie 4). Cone cell Ca^2+^ spikes were readily seen without an adjacent IOC wave, and conversely, many cone cells were silent when IOC waves passed by (two such examples are indicated by arrows at 28s). While all secondary and tertiary pigment cells participated in propagating the IOC waves, not all cone cells were active during the 5-min recording (Fig. 4B). Of the 141 clusters analyzed, only 51 (35.7%) had all four cells exhibiting Ca^2+^ transients (Fig. 4C), and tabulation of the distribution indicated that 124 of the 564 (22.0%) cone cells were inactive. Moreover, the cone cell Ca^2+^ spike characteristics were noticeably different from those of IOC waves (Fig. 4D). For instance, from this recording, Ca^2+^ spikes (n=280) from 39 randomly selected IOC cells had a T_on_ of 12.03 ± 8.37s and a T_off_ of 22.18 ± 8.45s. In contrast, cone cell Ca^2+^ spikes (n=6179) had a shorter duration, with a T_on_ of 2.36 ± 0.34s and a T_off_ of 3.50 ± 1.04s. These Ca^2+^ spike differences, along with the lack of apparent coordination, demonstrate that IOC waves and cone cell Ca^2+^ spikes are separate events.

**Figure 4.**
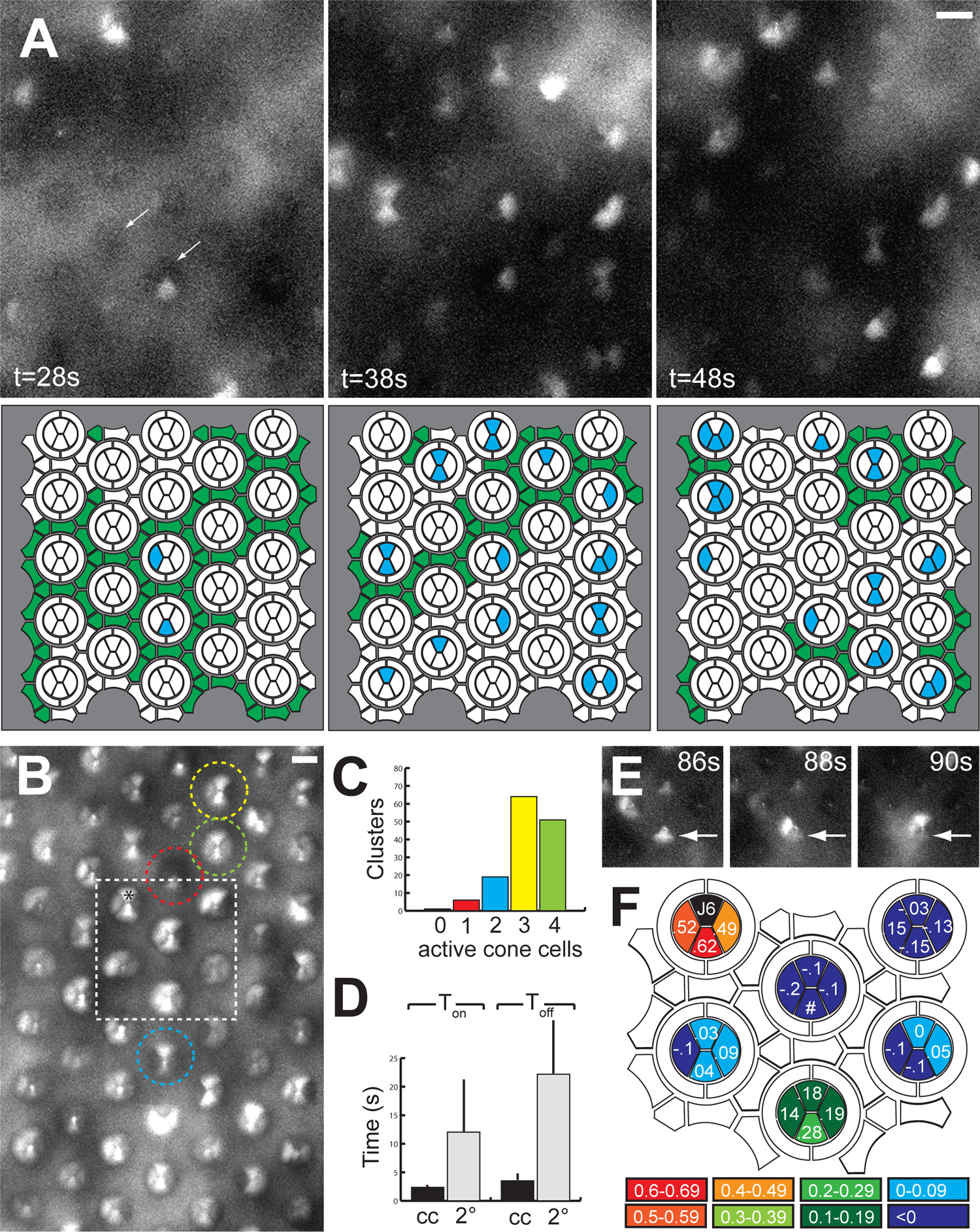
Cone cells exhibit Ca^2+^ activities distinct from the IOC waves. (A) Top row: a young *lGMR>GCaMP6m* retina showing Ca^2+^ activities in cone cells. Bottom row: IOC and cone cells experiencing Ca^2+^ spikes are highlighted in green and blue respectively. The secondary and tertiary pigment cells assigned to display Ca^2+^ spikes are approximations. Scale bar=10μm. (B) Maximum intensity projection of the same *lGMR>GCaMP6m* retina. Examples of clusters with one (red), two (blue), three (yellow), and four (green) *GCaMP6m*-active cone cells are labeled with dashed circles. Scale bar=10μm. Quantification of these clusters, with the same color code, is shown in (C). (D) The Ca^2+^ spikes in cone cells (n=564) have shorter T_on_ and T_off_ than the surrounding IOCs (n=39). (E) A cropped montage showing a Ca^2+^ spike, initiated at t=86s in an equatorial cone cell (arrow), sweeping through two cone cells of the same cluster. (F) A schematic map showing pair-wise Pearson correlation analysis performed on numeric series generated from GCaMP6m intensities over time with cell J6 (black; asterisk in B). Region containing analyzed clusters corresponds to the dashed box in B.

Interestingly, cone cells within a given cluster showed coordinated Ca^2+^ spikes. One example is shown in Fig. 4E, in which a cone cell Ca^2+^ spike initiated at t=86s (arrow) swept through the adjacent two cells in successive time frames. This suggests cone cells in each cluster form their own mini wave network, independent of the IOC waves and other clusters. In support of this, pair-wise Pearson analysis of GCaMP6m signal over time with cone cells of the same cluster showed higher correlations than with those from different clusters (Fig. 4F).

### IOC waves start at late pupal stage and persist in older flies

To understand how these Ca^2+^ wave networks are constructed, we characterized the onset and pattern of GCaMP6m signals in developing retinas. Ca^2+^ waves were observed in *ey>GCaMP6m* eye discs behind and ahead of the MF (Fig. S4; Movie 5). The eye disc is highly dynamic with multiple developmental events compressed in a strong temporal gradient (Roignant and Treisman, 2009, Tsachaki and Sprecher, 2012, Wolff and Ready, 1991). Accordingly, we focused on Ca^2+^activities in the latter half of pupal development, when pattern formation is complete (Cagan and Ready, 1989) and the temporal gradient has flattened; all cells are developing on the same timeline. At P9-P10 stages, *lGMR>GCaMP6m* retina showed no detectable Ca^2+^ activity (not shown). At P12 stage, while most of the cells were still silent, Ca^2+^ activities in isolated IOCs and cone cells began to emerge (Fig. 5A, Movie 6). Furthermore, instances of Ca^2+^ spikes propagating between adjacent IOC cells were seen (Fig. 5A and B), suggesting that the formation of IOC wave network has commenced. Imaging the same eye one day later showed widespread IOC waves and cone cell blinking (Fig. 5C, Movie 7), suggesting that the construction of the entire IOC wave network took place within this time.

**Figure 5.**
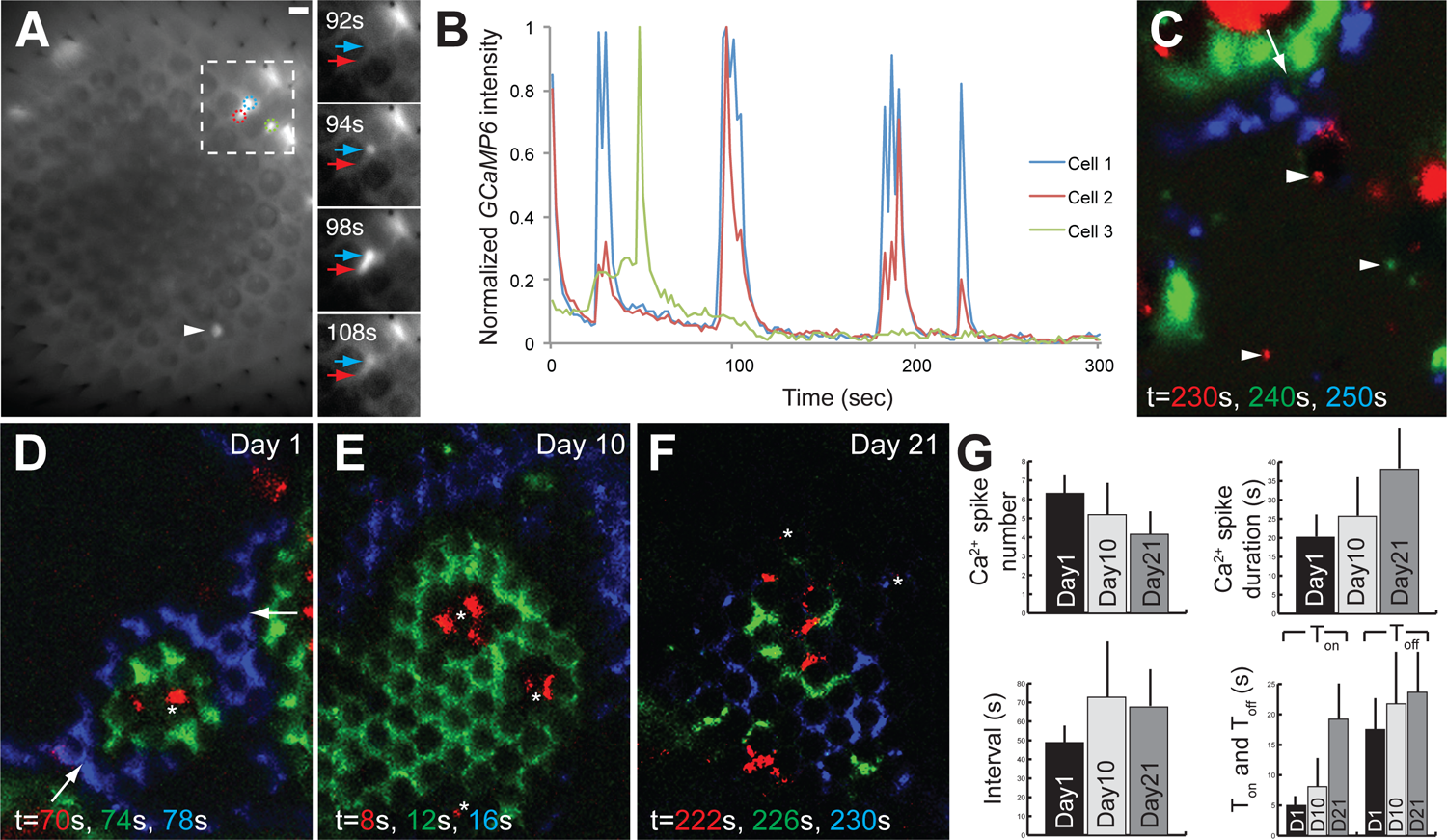
IOC waves emerge at late pupal stage. (A) Maximum intensity projection from a *lGMR>GCaMP6m* pupal eye at P12 stage. While most of the cells were inactive, a few IOC (color circles) and cone (arrowhead) cells exhibited Ca^2+^ transients. A region (dashed box) containing a two-cell “wave” is shown at a higher magnification. At t=94s, a tertiary pigment cell (blue arrow) experienced a Ca^2+^ increase, which was followed by a spike in an adjacent secondary pigment cell (red arrow) at t=98s. Scale bar=10μm. (B) Plot of normalized *GCaMP6m* intensity over time showing the Ca^2+^ oscillations in these two cells, but not the green cell, were coordinated. (C) An image overlaying three wave fronts from the same retina (shown in A) imaged one day later. An IOC wave, initiated at the dorsal edge at t=230s (red), was seen propagating ventrally. In addition, Ca^2+^ spikes in cone cells were readily seen (arrowheads). (D-F) Overlaying images showing wave fronts in 4s intervals from 1-day old (D), 10-days old (E), and 21-days old (F) *lGMR>GCaMP6m* retina. (G) Comparison of Ca^2+^ spike characteristics from IOC waves seen in *lGMR>GCaMP6m* retina of different age: 1-day (n=424; n=number of cells analyzed), 10-days (n=464) and 21-days (n=246).

To determine whether these Ca^2+^ events deteriorate with age, we analyzed *lGMR>GCaMP6m* retinas from older flies. Young retina, as abovementioned, showed regular IOC waves and cone cell blinking (Fig. 5D). Whereas cone cell Ca^2+^ spikes ceased about 5 days after eclosion, IOC waves were still present in 10- and 21-days-old *lGMR>GCaMP6m* retinas (Fig. 5E&F). However, compared to the waves in younger retina, IOC waves in older flies appeared irregular, evidenced by the decrease in peak number and the large variation in spike intervals (Fig. 5G).

### IOC waves are independent of phototransduction

As the pigment and cone cells surround the photoreceptors, we asked whether the abovementioned Ca^2+^ events require phototransduction. Prior to each recording session, dark adapted *lGMR>GCaMP6m* retina was illuminated with 647nm light pulses to inactivate the metarhodopsin in photoreceptors. Nonetheless, IOC waves were often present immediately at the start of recording, suggesting that waves were spontaneous and ongoing in the dark. Still, we frequently observed a surge in the overall eye brightness with multiple waves merging to cover the whole retina within the first 30s following illumination, suggestive of initial GCaMP6m signal being influenced by light. To definitively test whether the occurrence of IOC waves requires phototransduction, we monitored *lGMR>GCaMP6m* signals in retina mutant for *norpA*, the *Drosophila* phospholipase C β homolog known for its role in acting downstream of the rhodopsin (Bloomquist et al., 1988). We reasoned that if the initiation or propagation of IOC waves requires phototransduction, GCaMP6m signals should be absent or noticeably reduced in *norpA*. While eyes mutant for *norpA^36^*, a complete loss-of-function allele, displayed no response to light stimulation (Pearn et al., 1996), robust IOC waves were seen in *norpA^36^* (Fig. 6A, Movie 8) and the Ca^2+^ transient profiles were similar to those observed in wild type (Fig. 6B). These results demonstrate that IOC waves are independent of phototransduction.

**Figure 6.**
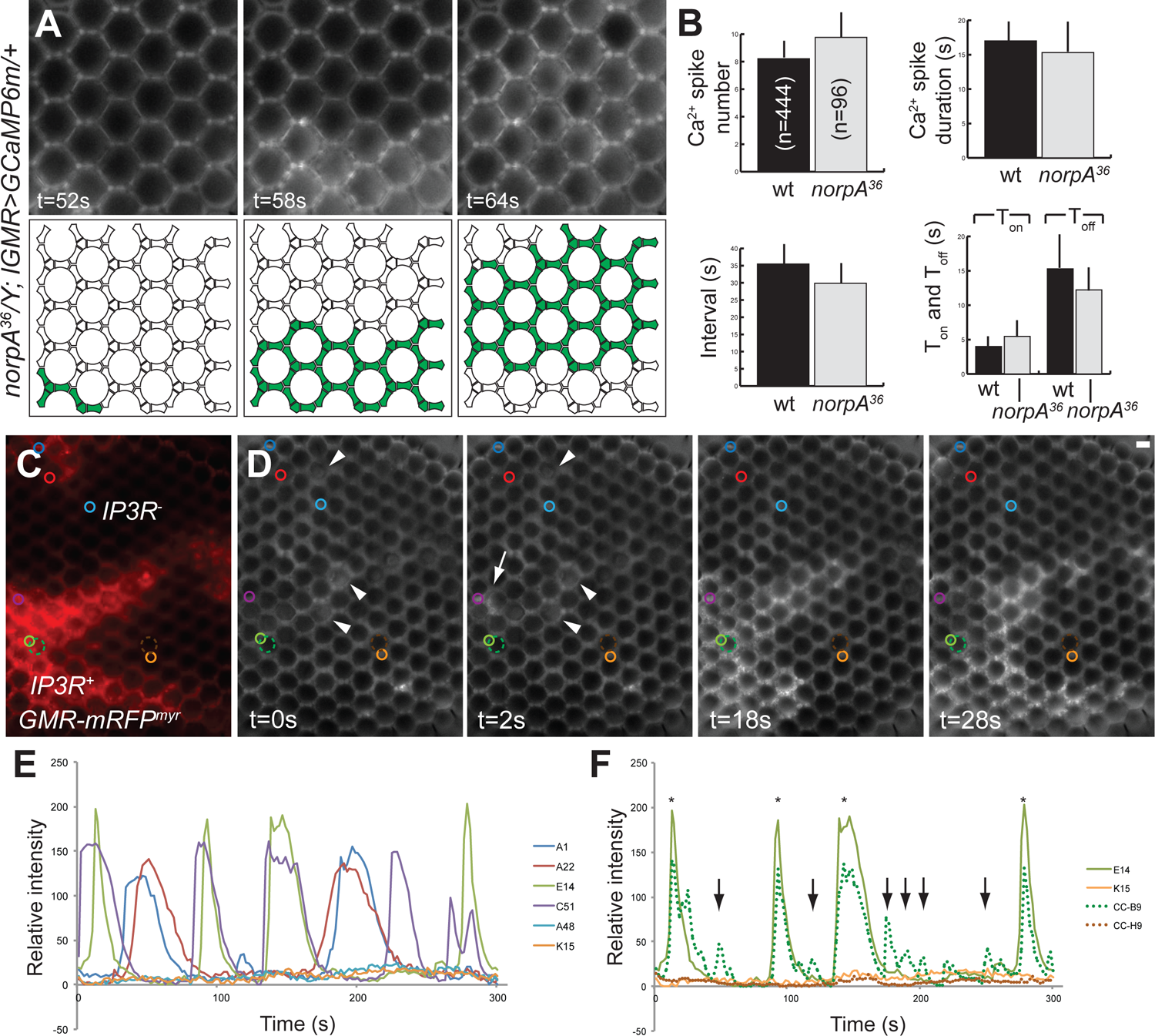
IOC waves are independent of phototransduction but require IP3R. (A) Top row: a montage showing an IOC wave moving across a young *norpA^36^* retina. Bottom row: schematic drawings showing the IOCs experiencing Ca^2+^ spikes (green). Scale bar=10μm. (B) Comparison of Ca^2+^ spike characteristics between wild type (wt; n=444) and *norpA^36^* IOCs (n=96). (C) A mosaic (*ey-FLP/+; lGMR>GCaMP6m/+; FRT^82B^, l(3)itpr^90B.0^/FRT^82B^, GMR-mRFP^myr^*) eye containing *IP3R^+^* and *IP3R^-^* patches, labeled by the presence and the absence of membrane-associated mRFP (myristylated mRFP) respectively. (D) A montage of GCaMP6m signals from the same eye. The IOCs were inactive at t=0s, and a Ca^2+^ wave initiated at the posterior edge at t=2s (arrow) propagated only in the IOCs within the large *IP3R^+^* clone. Ca^2+^ activities in cone cell clusters (arrowheads) could also be discerned. Scale bar=10μm. (E) Plot of *GCaMP6m* intensity *(F-F_min_)* over time in six IOCs (labeled by solid color circles in A) showing robust Ca^2+^ oscillation in the four *IP3R^+^* cells, but baseline in the two *IP3R^-^* cells. It is notable that the IOCs from distinct *IP3R^+^* clones (the red and blue cells from the smaller clone near the posterior-dorsal border, and the green and purple cells from the central large clone) exhibited different patterns of Ca^2+^ spikes. (F) Plot of *GCaMP6m* intensity over time in two cone cell clusters (labeled by dashed color circles in A; we quantified signal intensities from these cell clusters because the cone cells were out of focus and could not be individually identified). As IOC wave passage masks the *GCaMP6m* signals in cone cell clusters (asterisks), we compare the detection of Ca^2+^ spikes in *IP3R^+^* and *IP3R^-^* clusters during the absence of IOC wave (arrows). Intensity plots of two IOCs (green and orange solid circles; both are shown in C) adjacent to selected cone cell clusters are shown as reference.

### IOC waves and cone cell Ca^2+^ activities both require *IP3R*

Based on the fast speed (4-5um/s) of IOC waves (Jaffe, 2008), we speculated that the source for the IOC Ca^2+^ spikes is IP3R-regulated Ca^2+^ release from the ER stores. To test this, we monitored GCaMP6m signals in retinal cells mutant for *l(3)itpr^90B.0^*, a complete loss-of-function allele of *IP3R* (Venkatesh and Hasan, 1997). As *IP3R* is an essential gene, mosaic eyes containing clones of homozygous *l(3)itpr^90B.0^* cells, marked by the absence of a membrane-associated mRFP (Chang et al., 2002), were generated by FLP-induced recombination (Fig. 6C, Movie 9). Robust IOC waves were seen in wild type cells (*mRFP^+^*), but did not traverse into *l(3)itpr^90B.0^* mutant territory (Fig. 6D), indicating that IP3R function is required for wave propagation. Moreover, no GCaMP6m flash was observed in *l(3)itpr^90B.0^* cells, suggesting that IOC wave initiation, like propagation, requires IP3R function.

This mosaic eye contained two wild type patches partitioned by an *IP3R^-^* clone, allowing us to investigate the behavior of IOC waves from two separate populations in the same retina. As shown in Fig. 6E, IOC waves in the larger clone had more Ca^2+^ spikes (4.0 ± 1.2, n=210) than the waves in the small clone near the dorsal posterior region (2.4 ± 0.9, n=21). In addition, the Ca^2+^ spikes from these two clones occurred at different time points, indicating that the IOC waves could arise independently in separate cell populations.

At this focal plane, although the identity of specific cone cells experiencing Ca^2+^ spikes cannot be ascertained (out of focus), wild type clusters with cone cell Ca^2+^ activities were still discernable (Fig. 6D, arrowheads). In comparison, such clusters were completely absent in *IP3R^-^* cone cells (Fig. 6F), demonstrating that both IOC waves and cone cell Ca^2+^ activities require internal Ca^2+^ release.

### *IP3R* mutation disrupts IOC endfeet stress fiber morphology and retinal floor condensation

To understand the functional relevance of IOC waves, we looked for morphological defects associated with *IP3R^-^* IOCs. As removal of *IP3R* function has no effect on the retinal cell fate (Acharya et al., 1997, Raghu et al., 2000), we suspected that *IP3R^-^* mutation perturbs specific subcellular structures in IOCs. To ask whether *IP3R^-^* mutation affects the actin stress fibers at the endfeet, adult *l(3)itpr^90B.0^* mosaic eyes were stained with phalloidin to label the filamentous actin. These eyes were also rendered pigmentless with *cn* and *bw* mutations to eliminate interference from the pigment granules during imaging. In wild type clones, dense patches of fully contracted stress fibers surrounded and interconnected the grommets (arrows; Fig. 7A-C). In contrast, stress fibers in *l(3)itpr^90B.0^* mutant cells, marked by the absence of *ubi-mRFP^nls^*, exhibited various degrees of abnormalities. In some *IP3R^-^* cells, especially those in smaller clones (an example in Fig. 7E, asterisk), the stress fibers retained the patched morphology although their profiles were enlarged. In others, the defect was more severe, as the stress fibers, while still anchored to the grommets, appeared frayed, stretched, and unbundled (arrowheads, Fig. 7C).

**Figure 7.**
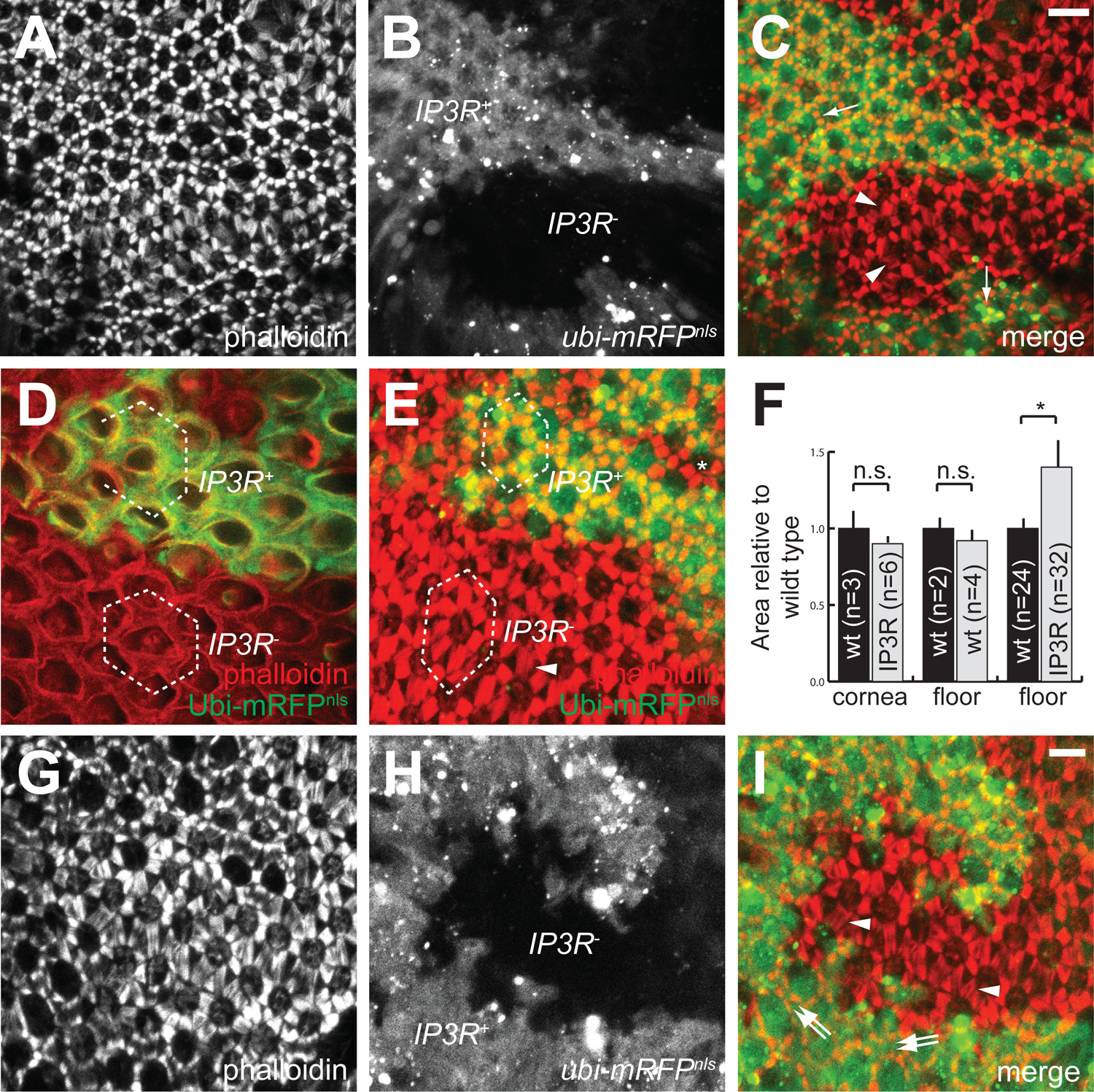
IOC endfeet stress fiber contraction and retinal floor condensation are disrupted in *IP3R* clones. (A-C) Maximum intensity projections of confocal sections (along the Z axis) of *ey-FLP; cn, bw; FRT^82B^, l(3)itpr^90B.0^/FRT^82B^, ubi-mRFP^nls^* adult eye stained with phalloidin (A; red in C). This *IP3R^-^* mosaic eye contains *IP3R^+^* and *IP3R^-^* patches, labeled by the presence and the absence of mRFP (B; green in C). Scale bar=10μm. (D-F) To measure the effect of *IP3R^-^* on the retinal floor area, hexagonal regions (D-E; dashed lines), generated by connecting the centers of six adjacent clusters, are quantified (F). Hexagonal regions at the cornea level (D) and in wild type clones at the floor level are included as controls. For each comparison, the area in *mRFP*-positive region is normalized to 1, and the numbers of hexagonal regions analyzed are indicated. n.s. not significant, *p<0.001(Student’s t test) as compared to internal control. (G-I) Maximum intensity projections of confocal sections of a mosaic retina at ∼75% pupal development stained with phalloidin (G; red in I). The *IP3R^+^* and *IP3R^-^* patches are labeled by the presence and the absence of mRFP (H; green in I). At this stage, the actin stress fibers within each *IP3R^+^* IOC have not yet consolidated into one single structure, thus appearing as two distinct patches abutting adjacent grommets (double arrows). In comparison, the stress fibers in *IP3R^-^* are frayed, appearing as filaments emanating from the grommets (arrowheads). Scale bar=10μm.

To ask whether this disruption of endfeet stress fibers affects the retinal floor area, we measured the cluster area, defined by a hexagon generated by connecting the centers from the 6 adjacent clusters (Fig. 7D-E). Mosaic eyes containing only wild type clones showed no difference in the size of these hexagons, demonstrating that FLP-induced mitotic recombination had no influence on the retinal floor area (Fig. 7F). In contrast, in mosaic eyes containing *IP3R^-^* clones, the retinal floor area per cluster in mutant patches was ∼40% larger than their sibling wild type clones. This perturbation of area size in *IP3R^-^* clones is specific to the retinal floor, as the apical area (the cornea level) in *IP3R^-^* clones remained unaffected.

### *IP3R* mutation affects endfeet stress fiber contraction

The aberrant stress fibers in adult *IP3R^-^* cells could be caused by a block in the progression of stress fiber contraction during pupal development or an inability to maintain the contracted fibers in adults. To distinguish between these possibilities, we monitored stress fiber morphology in pupal *IP3R^-^* mosaic eyes. If IP3R’s role is limited to maintaining contracted stress fibers in adults, stress fibers in *IP3R^-^* mutant pupal eyes should appear normal. In a *l(3)itpr^90B.0^* mosaic eye at ∼75% pupal development, the endfeet stress fibers in wild type cells were partially contracted, appearing as two discrete patches next to adjacent grommets (double arrows; Fig. 7G-I). In comparison, stress fibers in *IP3R^-^* mutant cells were frayed and unbundled (arrowheads), indicating that IP3R is required for the progression of endfeet stress fiber contraction. It is noteworthy that the stage in which this phenotype is manifested coincides well with the onset of IOC waves.

### *IP3R* loss perturbs basement membrane collagen IV

To investigate the effect of stress fiber deficits on the underlying ECM, we used vkg::GFP (Kelso et al., 2004), which fluorescently tags endogenous *Drosophila collagen IV α2* (Yasothornsrikul et al., 1997), a ubiquitous, network-forming basement membrane component (Kefalides, 1973, Brown et al., 2017, Fidler et al., 2017). In wild type adult retina, basement membrane vkg::GFP is organized in grommets and ridges (Fig. 1B, C). Compared to wild type, vkg::GFP signal intensity in *IP3R^-^* clones was reduced (Fig. 8A-D). Mean intensity measurement of vkg::GFP showed a 32% reduction in the abovementioned hexagons in *IP3R^-^* clones (wt n=2; *IP3R^-^* n=2; number of hexagons). These phenotypes were particularly noticeable in a mosaic cluster (dashed box; Fig. 8), in which the vkg::GFP intensity was lower on the *IP3R^-^* side with abnormal floor area and stress fibers. This selective area expansion on the *IP3R^-^* side also resulted in an apparent shift of grommet position towards the *IP3R^+^* side. Lastly, the grommet area in *IP3R^-^* clones was reduced by 25% (Fig. 8E; wt n=16; *IP3R^-^* n=29; n=number of grommets), suggesting that the stress fiber contraction provides a stretch to enlarge the grommet size. Taken together, these observations demonstrate that tensile forces from the stress fibers shape the basement membrane in connection with endfeet contraction.

**Figure 8.**
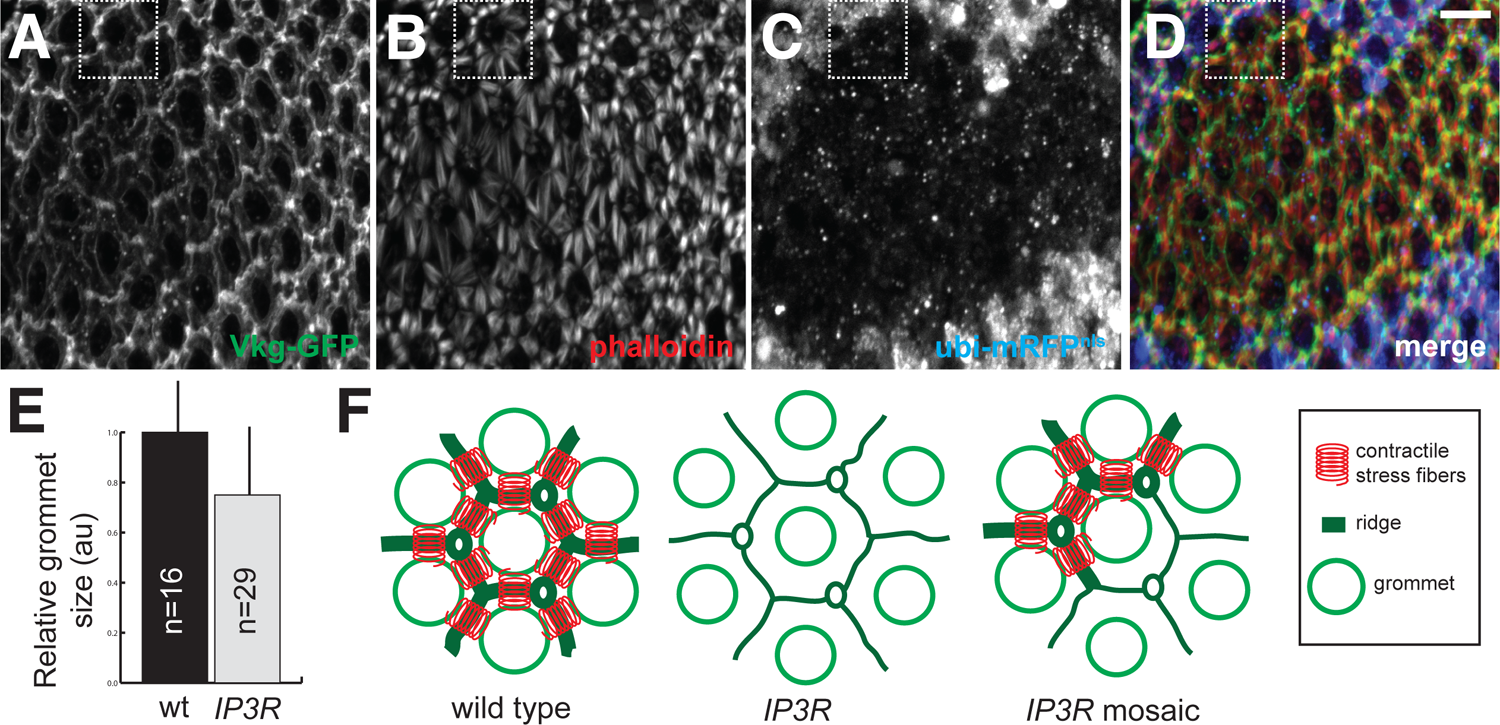
vkg::GFP morphology is altered in *IP3R* mutant. (A-D) Maximum intensity projections of confocal sections of an *ey-FLP; cn, bw, vkg::GFP; FRT^82B^, l(3)itpr^90B.0^/FRT^82B^, ubi-mRFP^nls^* eye stained with phalloidin (B; red in D). This mosaic eye expresses vkg::GFP (A, green in D) and contains *IP3R^+^* and *IP3R^-^* patches, labeled by the presence and the absence of mRFP (C; blue in D). A cluster containing a mixture of *IP3R^+^* and *IP3R^-^* cells is highlighted with a dashed box. Scale bar=10μm. (E) Quantification of grommet size in *IP3R^+^* and *IP3R^-^* patches. The numbers of grommets analyzed for each genotype are indicated, and the *p*-value from Student’s *t*-test is 0.003. (F) Schematic explanations of *vkg::GFP* morphology in various genotypes. In wild type, the pull by contracted stress fibers (red springs) enlarges the grommets and brings them closer together. In addition, the stress applied on ECM causes the materials between the grommets to crumple and form ridges. In contrast, as the contractile force is absent in *IP3R^-^* mutants, the grommets shrink in size and are farther apart, and the ECM ridges are diminished. In mosaic cluster, tensions are present on only one side of the cluster, resulting in an apparent shift of grommet position towards to the *IP3R^+^* side.

### Phosphorylated MRLC level is reduced at *IP3R* mutant retinal floor

To understand how IOC waves regulate the endfeet stress fiber contraction, we stained *l(3)itpr^90B.0^* mosaic eyes with a rabbit polyclonal antibody against phosphorylated MRLC (p-MLC, Ser-19). Ser-19 phosphorylation in MRLC, a target of MLCK in response to Ca^2+^ signaling, can augment the activity of non-muscle myosin II (Scholey et al., 1980, Heissler and Sellers, 2016). The p-MLC antibody detected signals in both wild type and *IP3R^-^* cells at the retinal floor, although p-MLC signal intensity was noticeably lower in *IP3R^-^* cells (Fig. 9). This result suggests IOC waves can impact stress fiber contraction by modulating the level of MRLC phosphorylation.

**Figure 9.**
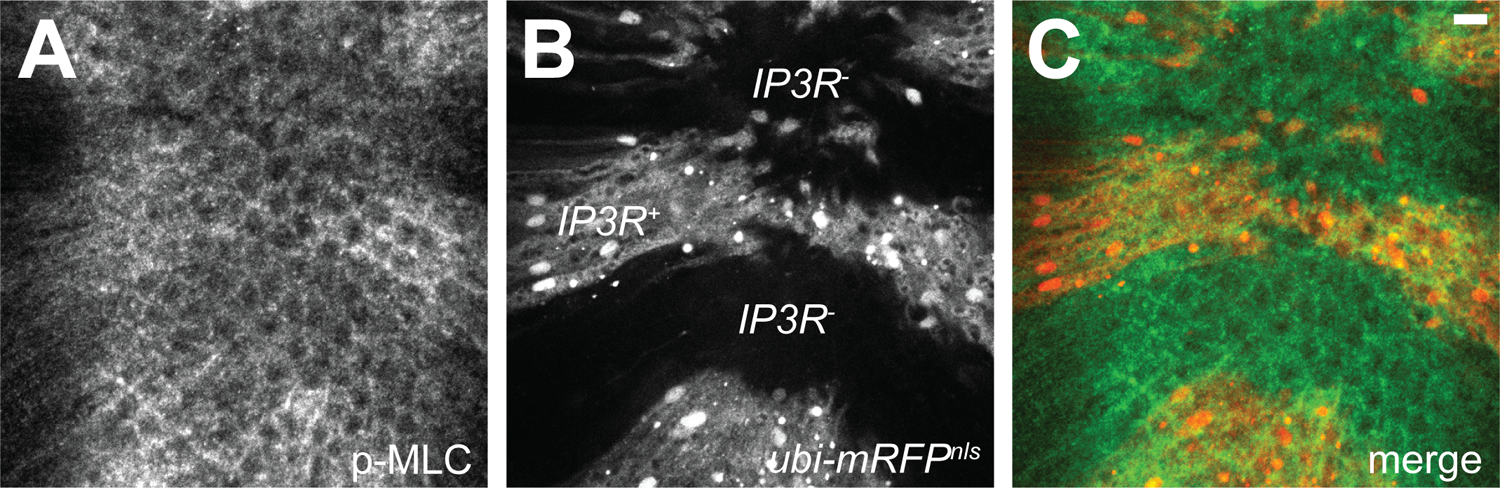
Phosphorylated myosin regulatory light chain is reduced in *IP3R^-^* cells. (A-C) Maximum intensity projections of confocal sections of *ey-FLP; cn, bw; FRT^82B^, l(3)itpr^90B.0^/FRT^82B^, ubi-mRFP^nls^* retina stained with anti-phospho-MLC (A; green in C). The *IP3R^+^* and *IP3R^-^* patches are labeled by the presence and the absence of mRFP (B; red in C). Scale bar=10μm.

## Discussion

Using *lGMR>GCaMP6m*, we have discovered the presence of intercellular Ca^2+^ waves that spontaneously and regularly sweep across the honeycomb lattice of IOC cells and the primary pigment cells. We provide evidence that this IOC-specific communication is insulated from the cone cell Ca^2+^ spikes and phototransduction but requires IP3R function. The IOC waves are formed during pupal development and persist with age. As intercellular propagation of Ca^2+^ spikes have been described in glia-glia networks in mammalian retina (Kurth-Nelson et al., 2009), our demonstration shows that communication with Ca^2+^ signaling is likely a fundamental and evolutionarily conserved feature of retinal accessory cells.

In adult *Drosophila* eye, photoreceptors depolarize with Ca^2+^ influx in response to light stimulation (Asteriti et al., 2017). As IOC and cone cells contact the photoreceptors, it seems plausible that the occurrence of IOC waves and cone cell spikes depends, at least in part, on phototransduction. The wave surges observed near the beginnings of recordings further lend support to the notion that IOC waves are influenced by illumination. However, the occurrence of IOC waves was clearly unaffected in a *norpA* null mutant. The characteristics of Ca^2+^ spikes in *norpA^36^* retina were comparable to those in wild type, suggesting that the complete removal of a phospholipase C critical for phototransduction has no modulatory effect on IOC waves. Moreover, while stimulus-independent Ca^2+^ oscillations in developing photoreceptor have recently been reported (Akin et al., 2019), these Ca^2+^ oscillations occur too early (between ∼55-95 hours after pupa formation) to influence the IOC waves (Akin et al., 2019), and their Ca^2+^ characteristics show no resemblance to those of IOC waves. These observations imply that IOC waves are insulated from photoreceptor activities. Likewise, a mechanism must exist to compartmentalize the IOC and cone cell Ca^2+^ networks. One possible mechanism may involve a restricted propagation of second messenger critical for Ca^2+^ increases. For instance, *Inx2* (*Innexin 2*), a *Drosophila* homolog of the gap junction component connexin, is expressed specifically in IOCs and localized to the endfeet (Richard and Hoch, 2015). As *Inx2* mediates the injury-induced slow Ca^2+^ waves in wing discs (Restrepo and Basler, 2016, Balaji et al., 2017), it may act similarly in the eye to regulate a cell-specific passage of second messenger required for IOC waves.

More importantly, using mutational analysis of *IP3R*, we have implicated this Ca^2+^ communication in facilitating and coordinating stress fiber contraction at IOC endfeet. During pupal development, the appearance of endfeet stress fibers transitions from arrays of filaments emanated from the grommets at mid-pupal stage to large patches at late pupal stage. (Baumann, 2004, Longley and Ready, 1995). The profiles of these structures continue to consolidate and appear as small dense patches by eclosion. In *IP3R^-^* adult and pupal mutant retinas, the morphology of stress fibers arrests at earlier stages. Concomitantly, the retinal floor in *IP3R^-^* clones increases by ∼40%, consistent with the notion that stress fiber contraction powers the retinal floor condensation. The apical surface in *IP3R^-^* clones remains unaffected, suggesting that *IP3R^-^* does not disrupt actin cytoskeleton globally and its effect on endfeet stress fibers is specific. Collectively, these results indicate that IP3R has a role in facilitating the progression of stress fiber contraction at the endfeet.

Several lines of evidence suggest IOC waves and endfeet stress fiber contractions are causally linked. First, the phenomena of IOC waves and endfeet compaction are spatially and temporally correlated. Both processes are manifested in the secondary and tertiary pigment cells, and the emergence of IOC waves overlaps with the duration of endfeet stress fiber contraction. In addition, elimination of IP3R function in the eye does not affect cell fate determination and phototransduction (Acharya et al., 1997, Raghu et al., 2000), suggesting these defects in IOC waves and stress fiber contractions are specific. Furthermore, MRLC Ser-19 phosphorylation at retinal floor in *IP3R^-^* clones is reduced, supporting a plausible mechanism in which IOC waves regulate myosin II activity through modulating MRLC phosphorylation. Based on these observations, we propose the removal of IP3R function abolishes Ca^2+^ increase from internal stores, which directly disrupts the occurrence of IOC waves. The absence of cytosolic Ca^2+^ increase in IOCs then inactivates myosin II at the endfeet, thereby inhibiting stress fiber contraction and retinal floor condensation. As Ca^2+^ increases in IOCs are elicited by waves, stress fiber contractions critical for retinal floor morphogenesis are coordinated.

The consistency of Ca^2+^ spikes in IOC waves makes this cellular communication well suited for coordinating endfeet stress fiber contraction for floor condensation. Analysis of IOC Ca^2+^ spikes suggest the presence of regular refractory period, which ensures all IOCs of the same eye experience a similar number of spikes, regardless of the frequency of IOC wave initiations. Furthermore, no increase in Ca^2+^ spike amplitude was seen in cells at wave merging, suggesting that all IOCs experience Ca^2+^ spikes of similar amplitude. Indeed, we confirmed that IOC Ca^2+^ spike parameters from different regions of the eye were similar. It is possible that this uniformity of Ca^2+^ spikes in IOC waves ensure the contractile forces are evenly applied across the retina during floor reduction. Ca^2+^ oscillations in cultured myofibroblasts have been shown to induce periodic micro-contractions of actin fibers (Castella et al., 2010), and it is likely that IOC waves have a similar effect on actin stress fibers at the pigment cell endfeet. In this scenario, regular passages of IOC waves would induce periodic and incremental contractions of the endfeet stress fibers to compact the retinal floor. Tensile forces driving the apical constriction of cells in the mesoderm furrow during gastrulation and pulling the epidermis cells to move dorsally during dorsal closure also occur in a pulsed fashion (Solon et al., 2009, Martin et al., 2009).

While the floor area increase in *IP3R^-^* implicates IOC waves in floor condensation, the phenotypic severity (40%) is significantly less than the fourfold area reduction required (Longley and Ready, 1995), indicating that IOC wave-mediated contraction is not the sole mechanism for basal surface compaction. In support of this, the pupal retinal floor reduction commences at ∼55% pupal development, earlier than the start of IOC wave formation. Thus, the retinal floor condensation consists of at least two phases: an early IOC wave-independent phase, which contributes to the majority of basement reduction, and a late IOC waves-dependent phase. We have observed mid-pupal retina containing clusters at different stages of floor reduction (Fig. S5D-F), implying that the condensation of ommatidial basement is initiated locally in patches and the clusters in different patches may compact at different rates. Tensile forces coordinated by IOC waves during the late phase may fine tune the floor condensation by smoothing out these differences from the early phase.

The retinal basement membrane, the ECM plane that defines the distal surface of the retina, is organized into two specializations, grommets and ridges. Grommets, the exit ports through which photoreceptor axons leave the retina, first form in early pupae at the completion of pattern formation (Fig. S5A-C). Ridges, dense linear ECM between adjacent grommets, emerge as stress fiber contraction draws grommets together. In wild type pupal eyes containing a mixture of fully and partially contracted stress fibers, ridges were prominent in regions with dense stress fibers (arrowheads, Fig. S5D-F). In *IP3R^-^* clones with reduced contraction, ridges are reduced. Together, these observations suggest ridges are raised as compression deforms and buckles the ECM meshwork. Ridges assemble into a hexagonal network oriented orthogonally to the contractile stress fiber network. As vkg/collagen IV stiffens ECM (Walma and Yamada, 2020), the ridge network may supply elastic energy that counterbalances contractile forces, thereby maintaining a stable retinal floor. The fly eye should be an excellent model for investigating how stress fiber contractile forces and ECM dynamics cooperate to shape tissue.

## Materials and Methods

### Drosophila genetics

All fly crosses were carried out at 25°C in standard laboratory conditions. *lGMR-GAL4* (8605), *UAS-GCaMP6m* (*20XUAS-IVS-GCaMP6m*; 42748), *norpA^36^* (9048), *FRT^82B^, GMR-src-mRFP* (7124), and *FRT^82B^, ubi-mRFP^nls^* (30555) were obtained from Bloomington Stock Center (Indiana, USA; the Bloomington numbers are indicated in parenthesis). Both *lGMR-GAL4* and *UAS-GCaMP6m* have transgenic insertions on the 2^nd^ and 3^rd^ chromosomes, and we have generated *lGMR>GCaMP6m* recombinants for respective chromosomes. Both *lGMR>GCaMP6m* recombinants exhibited similar Ca^2+^ waves; nonetheless, for simplicity and consistency, we used the 2^nd^ chromosome recombinant to generate all the movies presented. The *FRT^82B^, l(3)itpr^90B.0^* recombinant was a gift from Dr. Roger Hardie (Cambridge University, UK), and the *vkg::GFP* (*GFP^vkg-G00454^*) protein trap line (Morin, 2003) was a gift from Dr. Wu-Min Deng (Tulane University School of Medicine, USA).

To monitor Ca^2+^ activities in *IP3R* mitotic clones, *lGMR>GCaMP6m/CyO; FRT^82B^, l(3)itpr^90B.0^/TM3* males were mated with *ey-FLP/CyO; FRT^82B^, GMR-src-mRFP/TM3* females. *IP3R* clones in the adult eyes of *ey-FLP/lGMR>GCaMP6m; FRT^82B^, l(3)itpr^90B.0^/FRT^82B^, GMR-src-mRFP* progeny, easily recognized by the absence of membrane-associated mRFP, were imaged as described below. To examine actin stress fiber morphology in *IP3R* clones, *cn, bw; FRT^82B^, l(3)itpr^90B.0^/TM3* males were mated with *ey-FLP; cn, bw; FRT^82B^, ubi-mRFP^nls^/TM3* females. To monitor vkg::GFP localization in *IP3R* clones, *cn, bw, vkg::GFP; FRT^82B^, l(3)itpr^90B.0^/TM3* males were mated with *ey-FLP; cn, bw, vkg::GFP; FRT^82B^, ubi-mRFP^nls^/TM3* females.

### GCaMP6 imaging

For each Ca^2+^ activity recording, a live fly was glued eye-first onto a clean glass coverslip (Franceschini and Kirschfeld, 1971), maintained over an hour in the dark for adaptation, exposed to 647nm light pulses to shift active metarhodopsin to rhodopsin prior to imaging. Mounted flies were imaged on an inverted Nikon Eclipse TE2000 using a Nikon 40X 1.3 Plan Fluor objective. Time lapse movies were captured at one frame every 2 seconds, unless otherwise indicated, for 5 minutes using a Photometrics CoolSNAP camera controlled using Metamorph. Images were binned 2X2 and Nikon ND filters were used to attenuate excitation for typically 400ms exposures. No decrease in signal was observed over single or repeated observations. Mounting is non-injurious to eyes and animals; when kept in a humid chamber and fed with sugar water, it was possible to image flies repeatedly over several days.

### Imaging processing

The image files were rotated to bring anterior to the right and processed with ImageJ as described as below.

*a. <u>Sequential subtractions</u>*: Time-lapsed image stacks (2 sec per frame for a 5-minute recording) were separated into 151 individual png files with ImageJ (Image>Stacks>Stack to Images). A file series comprised of sequential subtractions of signal intensities from consecutive png files were generated in Python (code provided upon request). Composite files with color-coded wave fronts from various time points are generated with ImageJ (Image>Color>Merge Channels). This operation was only performed on IOC recordings because the Ca^2+^ spike intervals in the cone cell movie were too short.
*b. <u>Quantitative analysis of IOC GCaMP6m signals:</u>* To analyze the cellular signal intensities, the cells were labeled by hand in ImageJ with fixed-sized circular ROIs (5-pixel diameter), which were imported into ROI manager. Text files containing the mean signal intensities over time were then generated with the Multi-Measure function, and the numeric series were processed by a Python code (code provided upon request). The code normalized the intensity values from each cell by subtracting the minimal value in the series and divided by the range (i.e., F_normalized_ =(F-F_min_)/(F_max_-F_min_)). With an artificially cutoff of 0.24 (the value was empirically determined from applying the algorithm on several data sets, and F_normalized_ < 0.24 is considered noise), the number of peaks, the duration of peaks, the intervals between peaks, the time-onset for the peaks (T_on_), and the time-off for the peaks (T_off_) were counted.
*c. <u>Quantitative analysis of cone cell GCaMP6m signals:</u>* The cone cell movie had significant sample movements, which interfered with the extraction of mean intensity over time in individual cells. This issue was remedied by digitally stabilizing the image frames with the ImageJ Template-Matching plug-in. To improve the image quality, a minimum Z-stack projection was used as denominator for each frame with ImageJ Ratio Plus to make a ratio movie, followed by the ImageJ Enhance Contrast function. To characterize cone cell spikes, cells were manually labeled with fixed-sized circular ROIs and mean signal intensities over time were obtained with ImageJ as abovementioned. However, because the signal intensities in cone cells were dim and often influenced by the passages of IOC waves, the cone cell signal intensities were normalized by subtracting the signal intensity from a nearby IOC.
*d.* For comparing GCaMP6m in *IP3R^-^* mosaic eye, we obtained a mRFP map of the eye prior to the acquisition of time-lapsed stack of the GCaMP6m signal. A dual color movie of *IP3R^-^* mosaic eye was generated from 151 composite files, derived from merging individual GCaMP6m image files with the mRFP map file in ImageJ.

### Immunostaining and Western analysis

For immunostaining, dissected retinas were fixed with 4% paraformaldehyde and permeabilized with PBS + 0.3% Triton. For primary antibody, rabbit anti-phospho-MLC (3671T; SER-19; Cell Signaling Tech) polyclonal antibody was used at 1:10. Alexa-conjugated phalloidin and secondary antibodies were used at 1:100 and 1:100 dilutions, respectively. All images were acquired with Zeiss LSM 710 laser scanning confocal microscope. Zeiss Zen 2010 and Volocity (Improvision) softwares were used to render confocal stacks for 3D reconstruction.

## Acknowledgements

We thank Dr. Roger Hardie (Cambridge University, UK) for sharing the *FRT, IP3R* recombinant and discussion of IP3R phenotypes.

## Author contributions

D.F.R. and H.C.C. designed and performed the experiments, analyzed the results, and wrote the manuscript.

## Supplemental Figure Legends

**Figure 1. IOC Ca^2+^ transients are similar across the retina.** (A) A micrograph of *lGMR>GCaMP6/+* retina showing secondary (green) and tertiary (red) pigment cells labeled for quantitative analysis. A white line artificially demarcates the anterior and posterior halves. Scale bar=10μm. (B) Comparison of the Ca^2+^ peak number, interval, T_on_, and T_off_ between the secondary (n=204) and tertiary (m=69) pigment cells. (C) Comparison of Ca^2+^ spike parameters between IOCs in the anterior (ant; n=118) and posterior (post; n=155) halves. (D) A schematic map showing pair-wise Pearson correlation analysis performed on numeric series generated from GCaMP6m intensities over time with cell A1. Region containing analyzed clusters corresponds to the dashed box in A. The cells closest to A1 exhibit higher correlations than those farther away.

**Figure 2. Comparison of IOC wave Ca^2+^ spikes from two independent eyes.** (A) A micrograph montage of 10s intervals showing a Ca^2+^ wave initiated in the ventral region (dashed box) of a young *lGMR>GCaMP6m* retina at t=68s. Scale bar=10μm. (B) An image overlaying three wave fronts at t=68s (red), 78s (green), and 88s (blue) respectively. This IOC wave is the same one shown in A and corresponds to the green event in C. At t=78 sec, a distinct wave front (the yellow event in C) approaches from the anterior edge. (C) A schematic representation of the IOC wave origins during the 300s recording. Comparing to the movie shown in Fig. 2, this recording is noisier with more “flickering”, and only those resulted in waves are indicated. The color code for each event and its corresponding time is shown in the line below. (D) Plot of normalized *GCaMP6m* intensity (*(F-F_min_)/(F_max_-F_min_)*) over time showing the regular Ca^2+^ oscillation in 2 secondary pigments cells, selected from the dashed box in A. (E) Comparison of Ca^2+^ spike characteristics between the retina presented in Figure 2 (Eye 1; n=273) and this eye (Eye 2; n=474).

**Figure 3. IOC waves propagate through primary, secondary, and tertiary pigment cells.** (A) A young adult retina showing *GCaMP6m* signals from all three pigment cell types in different regions of the eye. Because of the retinal curvature, Ca^2+^ transients from secondary and tertiary pigment cells are readily recognized around the retinal periphery, whereas signals from the nuclei of anterior primary pigment cells in the central region are seen (arrows). In primary pigment cells, *GCaMP6m* signals in the nuclei are more prominent because of the quenching of cytoplasmic *GCaMP6m* fluorescence by pigment granules. The nuclei from the posterior primary pigment cells are not visible due to the angle of specimen preparation. At t=180s, wave initiations were seen at the anterior-ventral edge (arrowheads) and in nearby internal IOC cells (asterisk). At t=190s, another wave front was seen at the posterior edge (arrowheads). These waves then merged and moved dorsally across the *GCaMP6m* retina through all three pigment cell types. (B) Plot of normalized *GCaMP6m* intensity (*(F-F_min_)/(F_max_-F_min_)*) over time showing the Ca^2+^ oscillation in two primary (red and magenta lines), two secondary (blue and light blue), and two tertiary (green and light green) cells. The line colors correspond to color circles (selected cells) in A. The dip in *GCaMP6m* intensity around t=160s was caused by a sudden movement of the specimen. (C) Comparison of Ca^2+^ spike characteristics (spike duration, interval, T_on_, and T_off_) between primary (n=26), secondary (n=115), and tertiary (n=35) pigment cells.

**Figure 4. Ca^2+^ waves are present in larval eye discs.** (A) A micrograph montage of an *ey-GAL4; UAS-GCaMP6m* eye disc in 30s intervals, showing the presence of Ca^2+^ waves behind and ahead of the furrow (arrows). Instead of *lGMR-GAL4*, the larval eye disc was recorded with *ey-GAL4* to detect Ca^2+^ activities in cells anterior and posterior to the furrow. Ahead of the furrow, Ca^2+^ waves are present, but less frequent, and an example is shown initiating from the anterior edge at t=340s (blue arrow), reaching the furrow at ∼400s. At the furrow, when cells experience Ca^2+^ spikes, high GCaMP6m signals are seen in regularly spaced cell clusters (arrowheads). Posterior to the furrow, Ca^2+^ waves are frequent with higher GCaMP6m signals in pre-IOCs, resulting in a honeycomb-like appearance. Scale bar=40μm.

**Figure 5. The formation of grommets and ridges during pupal development.** A-F) Snapshots of 3D rendering of confocal sections of *cn, bw, vkg::GFP* pupal retinas stained with phalloidin (red). (A) Prior to rhabdomere extension (at ∼30% pupal development), unorganized vkg-positive structures are seen underneath the retina. (B) As the rhabdomeres (asterisks) are in the midst of extending throughout the depth of the retina (at ∼45% pupal development), vkg-positive cylindrical structures rise from the basement membrane to interface with the growing rhabdomeres. At this stage, the photoreceptor cell cortex, revealed by phalloidin staining, and vkg-positive cylinder assume the appearance of a wineglass (with photoreceptor cell cortex being the “bowl” and vkg cylinder being the “stem”). The junctures between the rhabdomeres and vkg::GFP stems constitute the future grommets (arrows). In addition, vkg-positive puncta are seen inside the IOCs (arrowheads), suggesting that these cells deposit the ECM into the basement membrane. (C) At the completion of rhabdomere extension (at ∼60% pupal development), elevated vkg::GFP signal labels the grommets (arrows, inset) and the IOC basal cell contacts at the retinal floor (arrowheads). A bottom view of a pupal eye at a similar stage as in (C) is shown in the inset. (D-F) At a later stage, the presence of vkg ridges correlates with the extent of actin stress fiber contraction, which varies across this retina. In regions where the stress fibers have contracted, the vkg ridges are prominent (arrowhead). In contrast, vkg ridges are reduced in regions where the stress fibers are extended (asterisk). Scale bar=10μm.

## Video Legends

All movies were generated at 10 frames/s from time lapse recordings captured at one frame every 2s.

**Movie 1.** Young *lGMR>GCaMP6m/+* adult retina at a distal plane (Fig. 2), revealing Ca^2+^ spikes in IOCs.

**Movie 2.** Young *lGMR>GCaMP6m/+* adult retina at a proximal plane (Fig. 3), showing Ca^2+^ spikes in primary pigment cells.

**Movie 3.** Young *lGMR>GCaMP6m/+* adult retina at a proximal plane (Fig. S3), showing Ca^2+^ spikes in primary pigment cells in the center and IOC waves in the periphery of the eye.

**Movie 4.** Young *lGMR>GCaMP6m/+* adult retina at a proximal plane (Fig. 4), showing Ca^2+^ activities in cone cells.

**Movie 5.** *ey>GCaMP6m/+* larval eye disc (Fig. S4). This time lapse was captured for a 10-minute duration.

**Movie 6.** *lGMR>GCaMP6m/+* pupal retina at approximately P12 stage (Fig. 5).

**Movie 7.** The same *lGMR>GCaMP6m/+* pupal retina from Movie 6 imaged one day later (Fig. 5).

**Movie 8.** Young *norpA^36^/Y; lGMR>GCaMP6m/+* adult retina at a distal plane (Fig 6).

**Movie 9.** Young *GMR>GCaMP6m/+* adult retina containing *IP3R^-^* mutant patches (Fig. 6).

